# Development of a Tick-Borne Encephalitis Virus Nanoparticle Vaccine Utilizing Envelope Dimer

**DOI:** 10.1101/2024.11.13.622924

**Authors:** Bingjie Wang, Haiyan Zhao

## Abstract

Tick-borne encephalitis virus (TBEV) is primarily transmitted to humans through tick bites, leading to symptoms of encephalitis with a fatality rate ranging from 1% to 30%, depending on the virus subtype. Currently, only inactivated virus vaccines are available for human use, though break-through infections can still occur. Therefore, developing new vaccines against TBEV is crucial. In this study, we designed and characterized a novel nanoparticle-based TBEV envelope (E) dimer vaccine. We successfully expressed and purified the TBEV E dimer by engineering disulfide bond mutants, and animal experiments demonstrated that the E dimer protein elicited a stronger immunogenic responses compared to the E monomer protein. Further, antibody depletion experiment confirmed that the E dimer protein effectively mimics the virion surface structure, inducing robust humoral immunity targeted at neutralizing epitopes. We also presented the TBEV E dimer on the surface of the nanoparticle Mi3 using the SpyCatcher-SpyTag system, with animal experiment showing that this TBEV E dimer nanoparticle vaccine elicited a potent humoral immune response. These findings offer new insights into the immunogenicity of the TBEV E dimer and suggest that the nanoparticle-based TBEV E dimer vaccine represents a promising and highly effective candidate for TBEV immunization.

## INTRODUCTION

Tick-borne encephalitis (TBE) is an infectious disease caused by the tick-borne encephalitis virus (TBEV), which leads to inflammatory symptoms in the central nervous system (CNS)[1]. TBE, which affects over 10,000 individuals annually worldwide, is a growing public health concern in Europe and Asia[2]. Humans are the definitive hosts of TBEV, which is primarily transmitted through tick bites and, less frequently, through the consumption of unpasteurized dairy products[1, 3]. Rare transmission routes include aerosol exposure in laboratory settings[4], blood transfusions[5], organ transplants[6], and vertical transmission from infected mothers to infants via breast milk[7]. TBEV is categorized into three primary subtypes: European (TBEV-Eu), Siberian (TBEV-Sib), and Far Eastern (TBEV-FE)[8, 9]. Recently, two additional subtypes have been identified: Baikal (TBEV-Bkl) and Himalayan (TBEV-Him)[10, 11]. The major TBEV subtypes differ in virulence, with TBEV-FE exhibiting the highest mortality rate (30%), compared to TBEV-Eu (1%-2%) and TBEV-Sib (6%-8%), which have lower mortality rates[9]. Infection with TBEV-Eu typically results in a biphasic disease course, whereas the Siberian and Far Eastern subtypes generally cause monophasic disease[12, 13]. Although the overall fatality rate is low, long-term neurological sequelae are common among survivors[1]. Notably, 40-50% of patients with acute TBE develop post-encephalitis syndrome (PES), which significantly impairs daily activities and quality of life. Common PES symptoms include cognitive impairment, altered sleep patterns, and neuropsychiatric issues such as apathy, irritability, and problems with memory and attention[8, 14, 15].

The tick-borne encephalitis virus (TBEV) is a member of the *Flaviviridae* family and the genus *Orthoflavivirus*, which also includes other arboviruses that infect humans, such as the yellow fever virus (YFV), dengue virus (DENV), Japanese encephalitis virus (JEV), Zika virus (ZIKV), and West Nile virus (WNV)[16]. TBEV is neurotropic, allowing it to infect the central nervous system (CNS) and resulting in various neurological pathologies; hence, the diseases caused by TBEV are collectively termed tick-borne encephalitis (TBE)[8]. Mature TBEV particles are approximately 50 nm in diameter and possess an envelope containing membrane (M) and envelope (E) proteins embedded in a lipid bilayer. The nucleocapsid consists of a capsid (C) protein and a single-stranded RNA genome, about 11,000 base pairs long. This RNA genome has an open reading frame (ORF) that encodes a polyprotein, which, after cleavage by viral and host proteases, is processed into three structural proteins (C, precursor M [prM], E) and seven non-structural proteins (NS1, NS2A, NS2B, NS3, NS4A, NS4B, NS5)[17, 18]. Non-structural proteins are crucial for viral replication, structural protein processing, and modulation of host cell functions. Glycoprotein E, the primary antigen of TBEV, is responsible for receptor binding and membrane fusion, making it the main target for vaccine development aimed at inducing a humoral immune response[17, 18]. The geographical distribution of TBEV spans Eurasia, from the Bordeaux region in France to Italy and Scandinavia in the south, and extends east to Siberia, China, and Japan[19]. This distribution is closely linked to the range of its tick vectors[3, 20, 21]. Most TBE cases are reported during the warmer months (April to November), coinciding with peak tick activity[14]. Human activities, such as deforestation and urbanization, influence climate and land use, creating favorable conditions for tick survival[22]. Consequently, new endemic areas have emerged, expanding TBE risk zones to previously unaffected regions and countries[23, 24], including the Netherlands, the United Kingdom, and higher alpine regions of Austria[25, 26, 27, 28].

The most important and effective protection against TBEV infection is immunization. Currently, all licensed TBEV vaccines are based on inactivated whole viruses, including various strains of the European or Far Eastern TBEV subtype[29, 30, 31]. These vaccines can generally be categorized into European, Russian, and Chinese types. Both European and Russian TBEV vaccines are considered safe and well-tolerated, with a proven efficacy in administration. However, both children and adults may experience mild to moderate systemic and local adverse reactions, such as fever, headache, or redness at the injection site[32, 33, 34, 35, 36]. Despite the widespread availability of vaccines in Europe, vaccine coverage remains relatively low, making TBEV control challenging[8, 37]. Additionally, immunization with the TBEV vaccine does not offer complete protection. Vaccine failures and breakthrough infections can occur in individuals with incomplete or irregular vaccination histories[38, 39]. To address these challenges, particularly in older adults, further research into optimized or novel vaccine strategies is necessary to ensure adequate protection for all at-risk groups[40]. Current developments in TBEV vaccine strategies include live attenuated vaccines and recombinant vaccines[41, 42]. Additionally, mRNA vaccines are being explored, wherein mRNA encoding the viral antigen is synthesized in vitro, encapsulated in lipid nanoparticles (LNPs), and then injected into the host. This approach induces the expression of the viral protein and elicits a robust immune response[43]. Notably, researchers have also developed mRNA-based vaccines for other flaviviruses, such as Powassan virus. A modified mRNA vaccine encapsulated in LNPs, encoding the Powassan virus prM and E genes, has demonstrated not only a robust immune response but also the induction of cross-neutralizing antibodies against various tick-borne flaviviruses[44, 45, 46].

In this study, we designed and characterized a TBEV nanoparticle vaccine displaying the TBEV E dimer. This vaccine elicited a strong humoral immune response, which was significantly better than that induced by the soluble TBEV E dimer protein. Moreover, compared with the monomeric E, the TBEV E dimer shows different antigenic sites, changes the surface structure, biases the immune response towards epitopes closer to the virus surface, thereby providing a powerful and effective neutralization effect. Our research results indicate that the TBEV E dimer is a promising immunogen for the development of TBEV vaccines, and the nanoparticle vaccine based on the TBEV E dimer is a potential vaccine candidate.

## RESULTS

### The V263C Mutation Can Effectively Promote the Formation of TBEV E Dimer

The E protein on the surface of the TBEV particle forms homologous dimers (Figure 1B), with a total of 90 dimers arranged to create an icosahedral shape. In previous studies involving DENV and ZIKV, researchers identified amino acids that interact closely at the dimer interfaces, one distributed between D3 of one E and D2 of another E, and the other distributed between D2 of one E and D2 of another E. These interacting amino acids were mutated to cysteine to promote the formation of disulfide bonds and stabilize the dimers[47, 48]. Following this approach, we investigated the dimer interface of the TBEV WH2012 E protein to identify the closest interacting amino acids. Since D3 of one E protein interacts with D2 of adjacent E at the fusion loop (FL), and this location corresponds to the highly conserved E-dimer epitope (EDE)[49], in order to prevent the introduction of mutations that destroy this epitope, we focused on introducing cysteine mutations between D2 of one E protein and D2 of another. Given the symmetric distribution of the dimer, it is theoretically possible to form a functional dimer by mutating just one amino acid.

**FIG 1.**
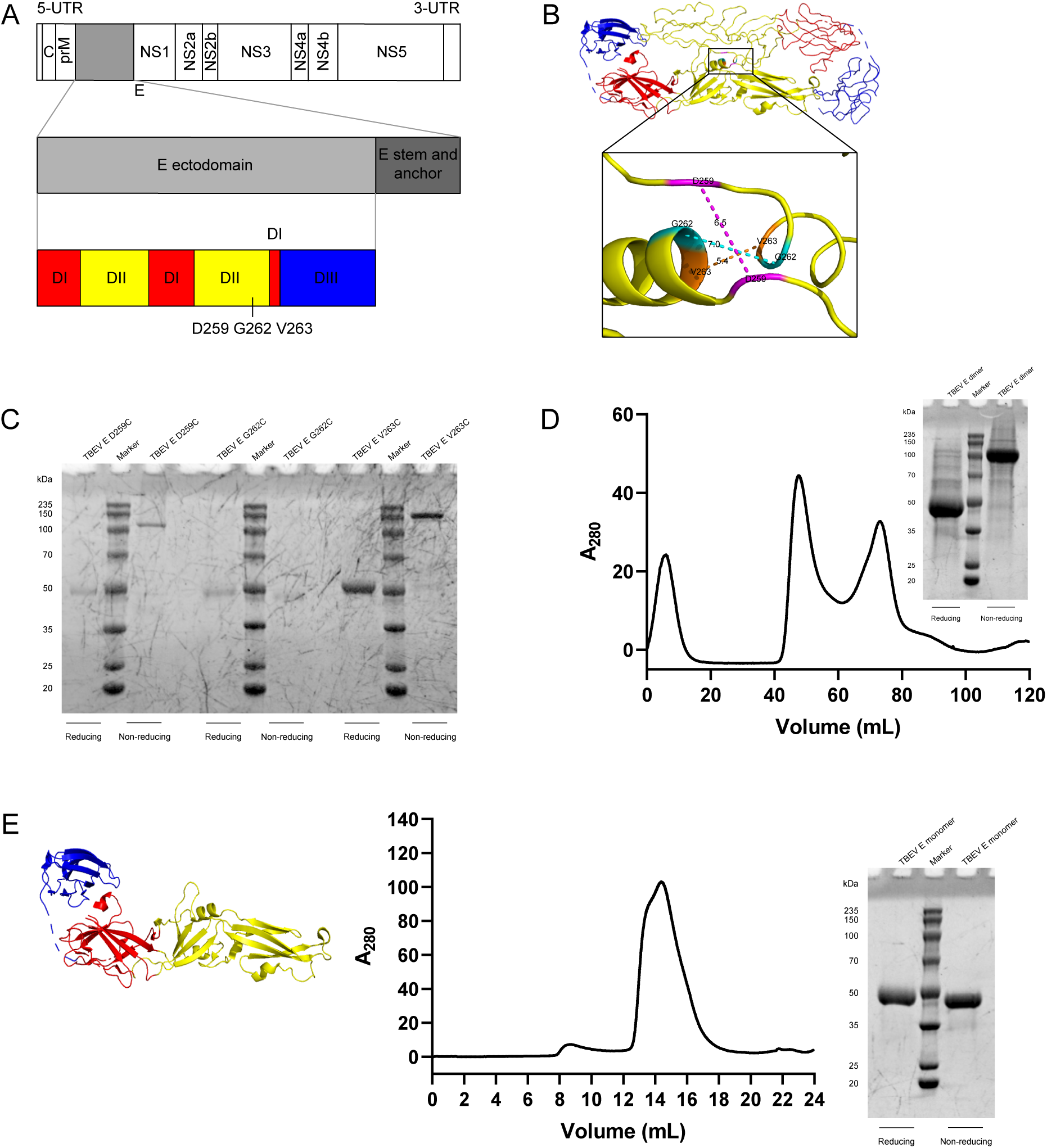
Expression and purification of TBEV E dimer and monomer proteins. (**A**) Schematic representation of the TBEV E protein in the genome.(**B**) Structure of the TBEV E dimer (PDB: 6J5G) highlighting the locations of mutations. The three mutations are D259C, G262C and V263C respectively.(**C**) The SDS-PAGE/Coomassie staining results of the mutant E proteins of D259C, G262C and V263C after nickel column purification.(**D**) Size-exclusion chromatography (SEC) and SDS-PAGE/Coomassie staining results following the expression of a large quantity of TBEV E V263C protein. The peak position of TBEV E V263C protein is approximately at 73 mL. The TBEV E V263C protein shows a monomer-sized band under reducing conditions and a dimer-sized band under non-reducing conditions.(**E**) Structural diagram, SEC, and SDS-PAGE/Coomassie staining results for the TBEV E monomer protein.

We identified three key amino acids at the dimer interface of the TBEV E protein: D259, G262, and V263 (Figure 1B). We mutated these residues to cysteine, constructed expression vectors, and performed protein expression and purification. SDS-PAGE/Coomassie staining of the purified proteins revealed that the G262C mutation did not effectively induce dimer formation, while both D259C and V263C mutations successfully produced dimers, with V263C showing more pronounced dimer bands, indicating a higher efficiency in dimer formation (Figure 1C). Therefore, we used the V263C mutant for subsequent experiments. Figure 1D illustrates the results of size-exclusion chromatography (SEC) and SDS-PAGE/Coomassie staining for the TBEV E V263C protein, showing a peak at approximately 73 mL, consistent with the theoretical dimer size. Under non-reducing conditions, distinct dimer bands were observed (Figure 1D). In comparison, we also expressed the TBEV E monomer protein, which showed only monomer bands under non-reducing conditions (Figure 1E).

### TBEV E Dimer Can Induce a Humoral Immune Response with a Stronger Neutralizing Ability

To compare the immunogenicity of the TBEV E dimer and monomer proteins, mouse immunization experiments were conducted. Each mouse was immunized three times with 10 *µ*g of TBEV E dimer or monomer protein intramuscularly with a three-week interval. Blood was collected from the orbital veins for ELISA and Reporter virus particle (RVP) neutralization experiments. Reporter virus particle (RVP) is prepared by co-transfecting pWNVI-GFP-replicon and the structural protein CprME of flavivirus into 293T cells (Figure S2)[50]. Additionally, we found that transfection of the structural protein of DENV could not effectively package RVP. Some researchers have found that some strains of Dengue virus type 2 (DENV2) are incompatible with the West Nile virus replicon protein, which is manifested in the low cleavage efficiency of the NS2B/3 protease of West Nile virus on the Dengue virus C protein[51]. A major obstacle to RVP assembly may be that the C protein cannot be fully released from its anchor peptide in the ER membrane. Therefore, we replaced the C protein of DENV with the C protein of TBEV and successfully achieved effective packaging (Figure S1).

The ELISA results against TBEV E dimer showed no significant difference in the IgG titers between the immune sera of TBEV E dimer and monomer proteins. Both titers increased significantly after the second immunization and remained stable thereafter (Figure 2B, S3). However, in the RVP neutralization experiment, the increase in IgG titer induced by the TBEV E dimer was accompanied by a significant enhancement in neutralization strength. In contrast, the monomer protein initially exhibited minimal neutralization ability, which gradually increased but never reached the level of the TBEV E dimer (Figure 2C, S4). These findings suggest that TBEV E dimer more accurately mimics the distribution of epitopes on the virus surface, resulting in a more focused and effective humoral immune response, and thus demonstrating superior neutralization ability.

**FIG 2.**
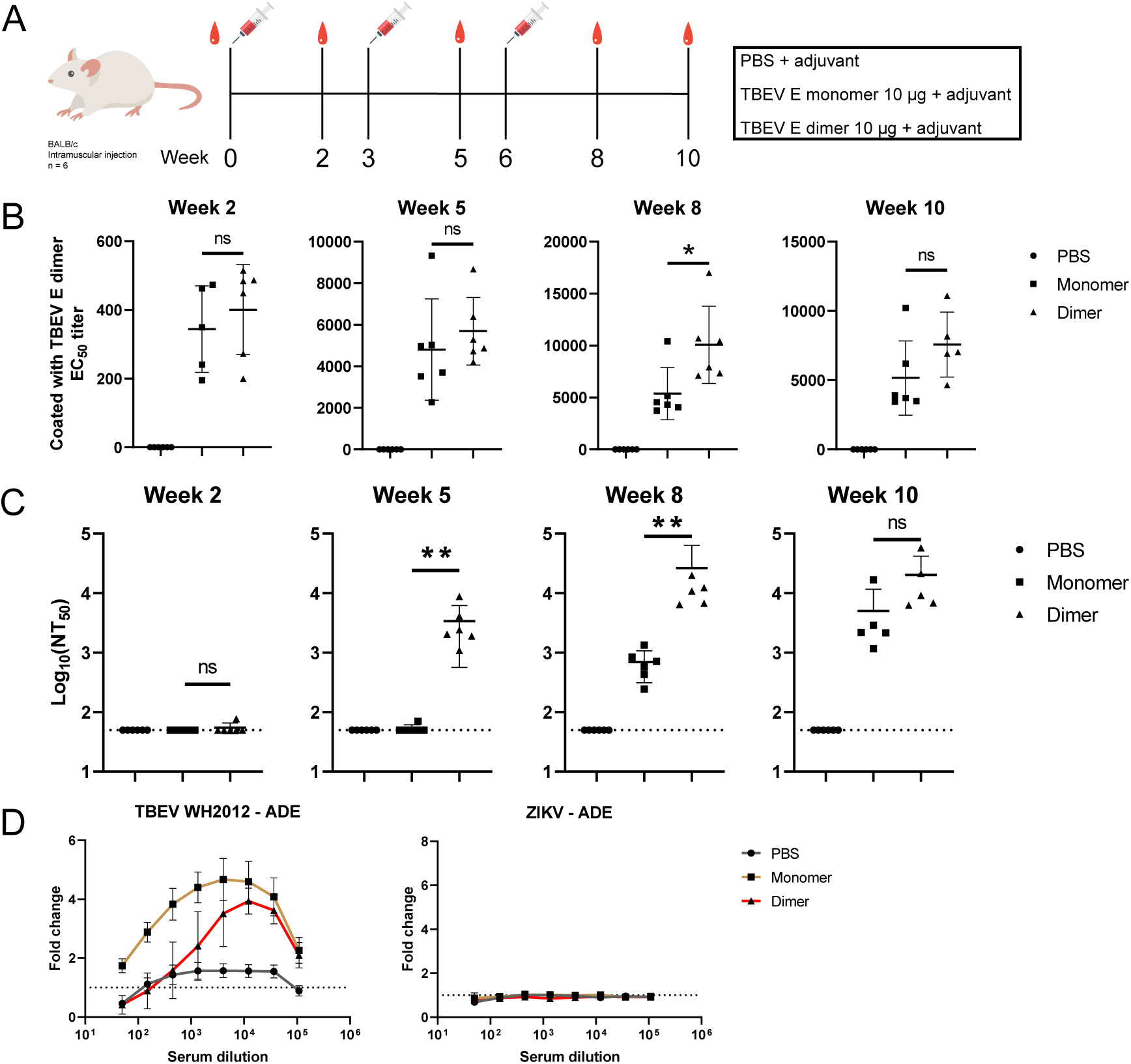
Evaluation of the immunogenicity of TBEV E dimer. (**A**) The mouse immunization schedule for the dimer and monomer proteins of TBEV E; (**B**) The IgG titers of sera binding to the TBEV E dimer after immunization with the TBEV E dimer and monomer proteins were determined by enzyme-linked immunosorbent assay (ELISA); (**C**) The neutralization activities of the sera after immunization with the TBEV E dimer and monomer proteins were determined by the reporter virus particle neutralization assay; (**D**) Using the mouse sera collected in the 8th week, detect the antibody-dependent enhancement of infection effects on TBEV WH2012 and ZIKV. ELISA, the reporter virus particle neutralization assay and the antibody-dependent enhancement of infection assay were performed independently twice in duplicate wells. The representative data presented, as well as the half-maximal effective concentrations (EC50) for ELISA and the half-maximal inhibitory concentrations (NT50) for neutralization, were determined using nonlinear regression analysis. The dashed lines indicate the limit of detection (LOD). Values lower than the LOD were calculated as the LOD. The data in each group were presented as the geometric mean and geometric standard deviation in one independent experiment. Statistical significance was determined by one-way ANOVA. ns indicates no significant difference, *p < 0.05, **p < 0.01, ***p < 0.001, ****p < 0.0001.

Currently, nearly all antibodies against flaviviruses induce antibody-dependent enhancement (ADE) of infection with their corresponding homologous viruses at sub-neutralizing concentrations or heterogeneous virus infection[52]. For instance, DENV has four serotypes; infection with one serotype followed by another can lead to ADE, exacerbating the disease and potentially leading to severe outcomes. Additionally, previous studies have indicated that DENV infection can enhance ZIKV infection through ADE[53]. To investigate the ADE effect, we tested the inhibitory activity of immunized serum in K562 cell which is widely used for ADE study for flavivirus. Both the E monomer and dimer proteins exhibited ADE effects on RVP - TBEV WH2012; however, compared with the immune serum of E monomer protein, the immune serum of E dimer protein observed the highest antibody-dependent enhancement (ADE) effect at a lower serum dilution (Figure 2D, S5). Furthermore, the immune sera of both the E monomer and dimer proteins showed no antibody-dependent enhancement (ADE) effect on Zika virus (ZIKV) in vitro, but showed an ADE effect on Powassan virus (POWV), and there was no difference between the two (Figure 2D, S6).

### The TBEV E Dimer Forms an ED12 Structure that the E Monomer Protein does not Have, which Can Induce Humoral Immunity with Strong Neutralizing Capacity

To further compare the humoral immunity induced by TBEV E dimer and monomeric proteins, we expressed various TBEV E domain proteins: ED123, ED12, and ED3. Previous studies have shown that the mutations T76R, M77E, W101R, and L107R jointly disrupt the epitope of the fusion loop (FL), preventing the binding of antibodies against FL[54]. Therefore, we also expressed ED123 T76R M77E W101R L107R (TBEV ED123 4mFL) (Figure S7). Additionally, we expressed ZIKV ED12 to evaluate cross-binding of humoral immunity. ELISA experiments revealed that both TBEV E dimer and monomeric protein-induced sera bind to TBEV ED123, ED12, and ED3 (Figure S8). Among these, the antibody titer against ED123 was the highest, followed by ED3, which was slightly higher than ED12 (Figure 3A). The FL mutations did not significantly change the antibody titer, indicating that the proportion of antibodies against FL in the serum is minimal (Figure 3B, S9). Both TBEV E dimer and monomeric protein-induced sera did not cross-bind with ZIKV ED12 (Figure 3A, S8).

**FIG 3.**
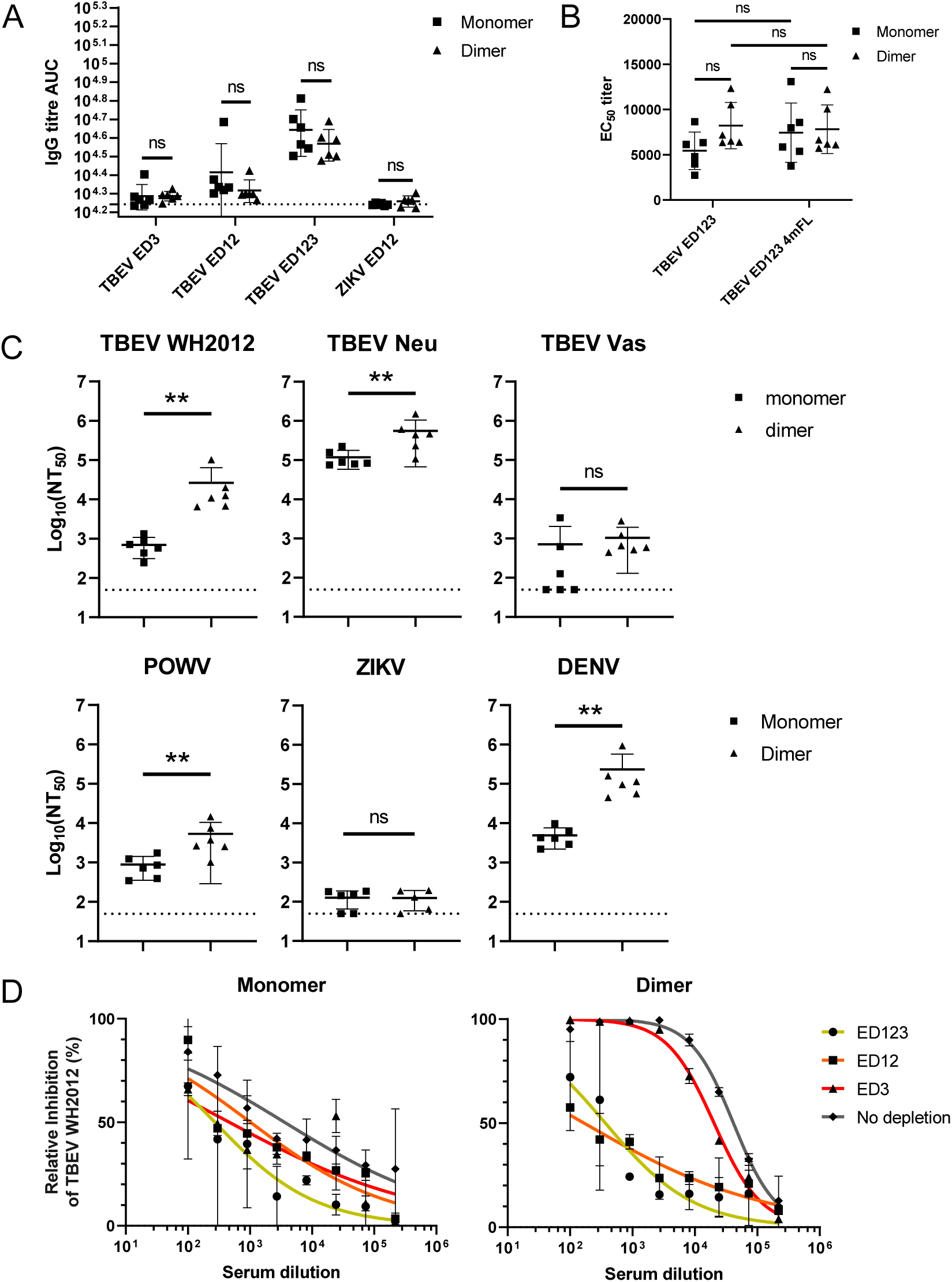
In-depth description of the immune response of TBEV E dimer. (**A**) The IgG titers of sera binding to TBEV ED3, ED12, ED123 and ZIKV ED12 of the mouse sera collected in the 5th week were determined by enzyme-linked immunosorbent assay (ELISA). The dashed line indicates the limit of detection (LOD) of the area under the curve (AUC).(**B**) The IgG titers of sera binding to TBEV ED123 and TBEV ED123 T76R M77E W101R L107R (TBEV ED123 4mFL) of the mouse sera collected in the 5th week were determined by enzyme-linked immunosorbent assay (ELISA).(**C**) The neutralization activities of the mouse sera collected in the 8th week against TBEV WH2012, TBEV Neu, TBEV Vas, POWV, ZIKV and DENV were determined by the reporter virus particle neutralization assay. Values lower than the LOD were calculated as the LOD. (**D**) In the antibody depletion experiment, the mouse sera collected in the 10th week were treated with TBEV ED123, ED12 and ED3 respectively, and then their neutralization activities against RVP TBEV WH2012 were determined. ELISA and the reporter virus particle neutralization assay were performed independently twice in duplicate wells. The representative data presented, as well as the half-maximal effective concentration (EC50) for ELISA and the half-maximal inhibitory concentration (NT50) for neutralization, were determined using nonlinear regression analysis. The antibody depletion experiment was also performed in duplicate twice. The data in each group were presented as the geometric mean and geometric standard deviation in one independent experiment. Statistical significance was determined by one-way ANOVA and two-way ANOVA. ns indicates no significant difference, *p < 0.05, **p < 0.01, ***p < 0.001, ****p < 0.0001.

We also compared the cross-neutralization effects of sera induced by TBEV E dimer and monomeric proteins against other subtypes of TBEV, POWV, ZIKV, and DENV. Notably, both TBEV E dimer and monomer protein-induced sera exhibited stronger neutralization effects against TBEV Neu (European subtype) (Figure 3C, S10). Previous studies have shown that the European subtype of TBEV has the lowest virulence, whereas the Far East subtype has the highest. These findings suggest that the virulence of different subtypes may be related to the host’s humoral immunity neutralization capacity. Consistent with the ELISA results, neither TBEV E dimer nor monomer protein-induced sera cross-neutralized ZIKV. However, TBEV E dimer and monomer protein-induced sera moderately neutralized TBEV Vas (Siberian subtype). The neutralizing ability of the immune serum of TBEV E dimer against RVP-TBEV WH2012, RVP-TBEV Neu and RVP-POWV was significantly stronger than that of the immune serum of the monomer protein, further illustrating the superior immunogenicity of TBEV E dimer.

To determine which domain contribute most to the neutralization capacity of the serum, we performed an antibody depletion experiment. Briefly, we added Strep tag to TBEV ED123, ED12 and ED3 proteins, bound them to streptavidin beads, and incubated them with sera induced by TBEV E dimer and monomer proteins. Antibodies bound to these proteins were removed with the beads, effectively depleting them from the serum. ELISA results indicated that these antibodies were successfully removed from both the TBEV E dimer and monomeric proteins (Figure S11). The RVP neutralization test conducted on the depleted serum showed that in the serum group induced by TBEV E dimer protein, the serum neutralization capacity significantly decreased after the depletion of antibodies against ED123 and ED12, but remained good after the depletion of antibodies against ED3 (Figure 3D). This indicates that the antibodies against the ED12 epitope contribute significantly to the neutralizing capacity of the serum after TBEV E dimer immunization, and ED3 also contributes to the neutralizing capacity, but relatively less. These results partially explain the previously observed phenomenon that although the humoral immunity induced by TBEV E dimer and monomer proteins shows similar antibody titers, TBEV E dimer shows a stronger neutralizing capacity. Specifically, antibodies against the ED12 epitope account for a large part of the neutralizing capacity induced by TBEV E dimer. The alteration of the antigen epitope distribution by TBEV E dimer forms epitopes with stronger neutralizing capacity against ED12, and these epitopes against ED12 cannot be formed in the E monomer protein.

### The TBEV E Dimer Can be Assembled with Mi3 in Vitro to Form Nanoparticle Vaccine

Compared to soluble protein antigen subunit vaccines, nanoparticle vaccines often induce a more robust humoral immune response[55]. For example, nanoparticle vaccines that utilize Mi3 nanoparticles assembled with various strains of RBD have demonstrated strong cross-neutralization capabilities[56, 57]. To enhance the humoral immune response further, we aimed to develop a nanoparticle vaccine incorporating the TBEV E dimer. Utilizing the SpyCatcher-SpyTag system[58, 59], which forms isopeptide bonds, we tagged the C-terminal of the TBEV E dimer with SpyTag. We then purified both Mi3 and the TBEV E dimer separately before assembling them in vitro (Figure 4A, 4B).

**FIG 4.**
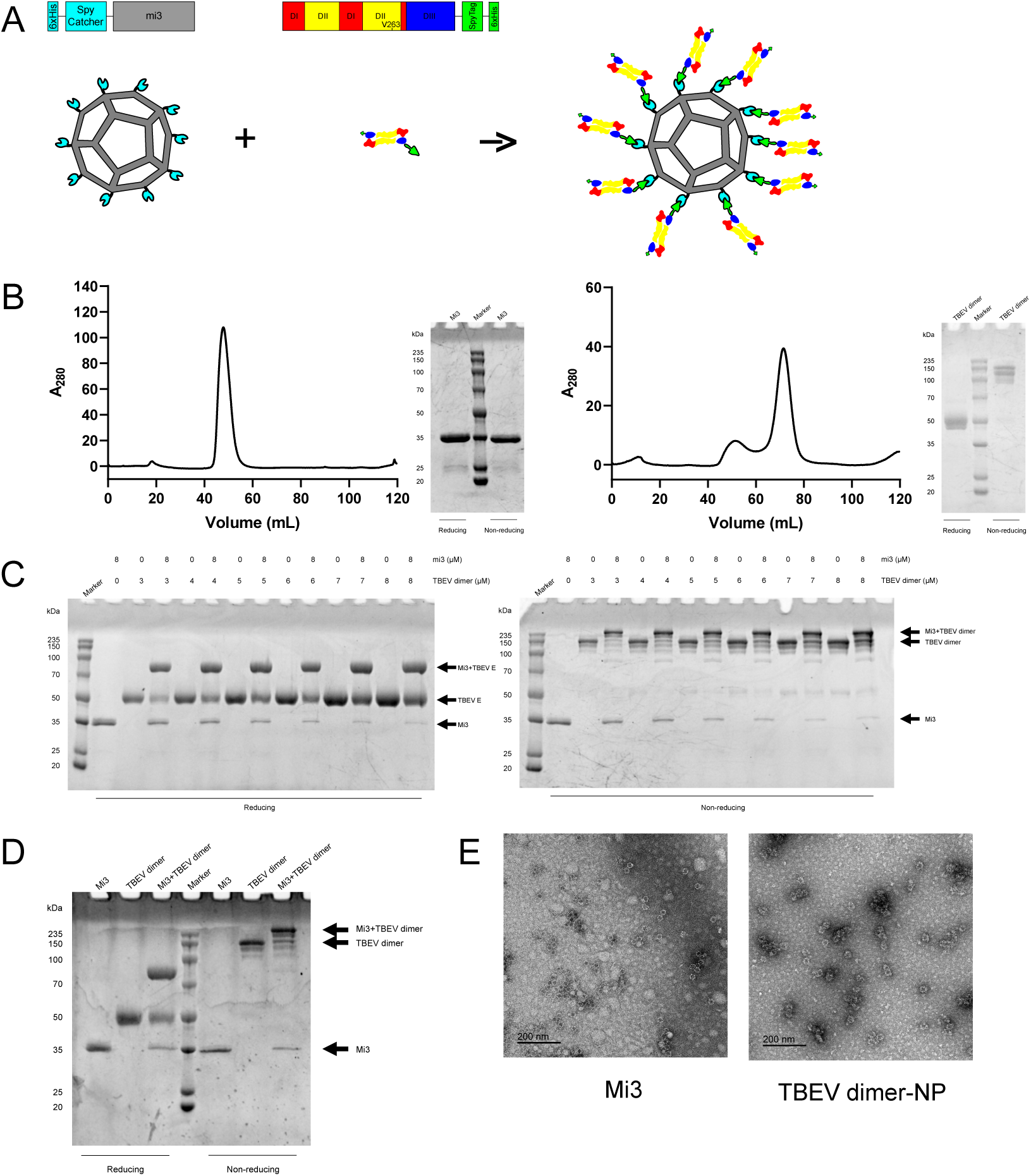
Characterization of the TBEV E Dimer Nanoparticle Vaccine. (**A**) Structural diagram of TBEV E dimer nanoparticle and their components.(**B**) The size exclusion chromatography (SEC) and SDS-PAGE/Coomassie staining results of Mi3 and TBEV E dimer - SpyTag. On the left is Mi3, and on the right is TBEV E dimer - SpyTag. (**C**) The SDS-PAGE/Coomassie staining results of Mi3 and TBEV E dimer - SpyTag at different assembly concentration ratios under reducing and non-reducing conditions.(**D**) The SDS-PAGE/Coomassie staining result after the assembly of 8 *µ*M Mi3 and 5 *µ*M TBEV E dimer - SpyTag before mice immunization. (**E**) Transmission electron microscopy images of TBEV E dimer nanoparticles and Mi3.

To determine the optimal assembly ratio, we mixed TBEV E dimer at various concentrations (3 *µ*M to 8 *µ*M) with a fixed concentration of Mi3 (8 *µ*M). We found that a 5 *µ*M concentration of TBEV E dimer mixed with 8 *µ*M Mi3 resulted in a nanoparticle assembly where the TBEV E dimer band in the free unassembled state was comparable to that at lower concentrations, and the free unassembled Mi3 band was comparable to that at higher concentrations. This indicates that the optimal assembly is achieved under these conditions, minimizing the presence of free TBEV E dimer and Mi3 (Figure 4C). Figure 4D displays the SDS-PAGE/Coomassie staining of the sample before immunization, based on the explored assembly condition. Additionally, transmission electron microscopy revealed that, compared to Mi3 nanoparticles, the assembled nanoparticles exhibited rougher and more irregular edges, indicating the successful assembly of TBEV E dimer on the surface of Mi3 (Figure 4E).

### Displaying TBEV E Dimer on the Surface of Nanoparticles Can Further Improve the Immunogenicity

We immunized mice twice with the prepared nanoparticle vaccine, 0.1 or 1 *µ*g/dose each time, at an interval of 3 weeks. Blood samples were collected from the orbital veins twice, and then the serum was used for ELISA and RVP neutralization experiments (Figure 5A). In the ELISA experiment, the IgG titer induced by the nanoparticle vaccine at 3 weeks was significantly higher than that of TBEV E dimer, indicating that preparing TBEV E dimer in the form of nanoparticles for immunization can induce faster and stronger humoral immunity (Figure 5B, S12). Consistent with the ELISA results, the serum immunized with the nanoparticle vaccine at 3 weeks had a stronger neutralizing capacity than the serum of TBEV E dimer (Figure 5C, S13). In conclusion, nanoparticle vaccines can further produce better humoral immunity.

**FIG 5.**
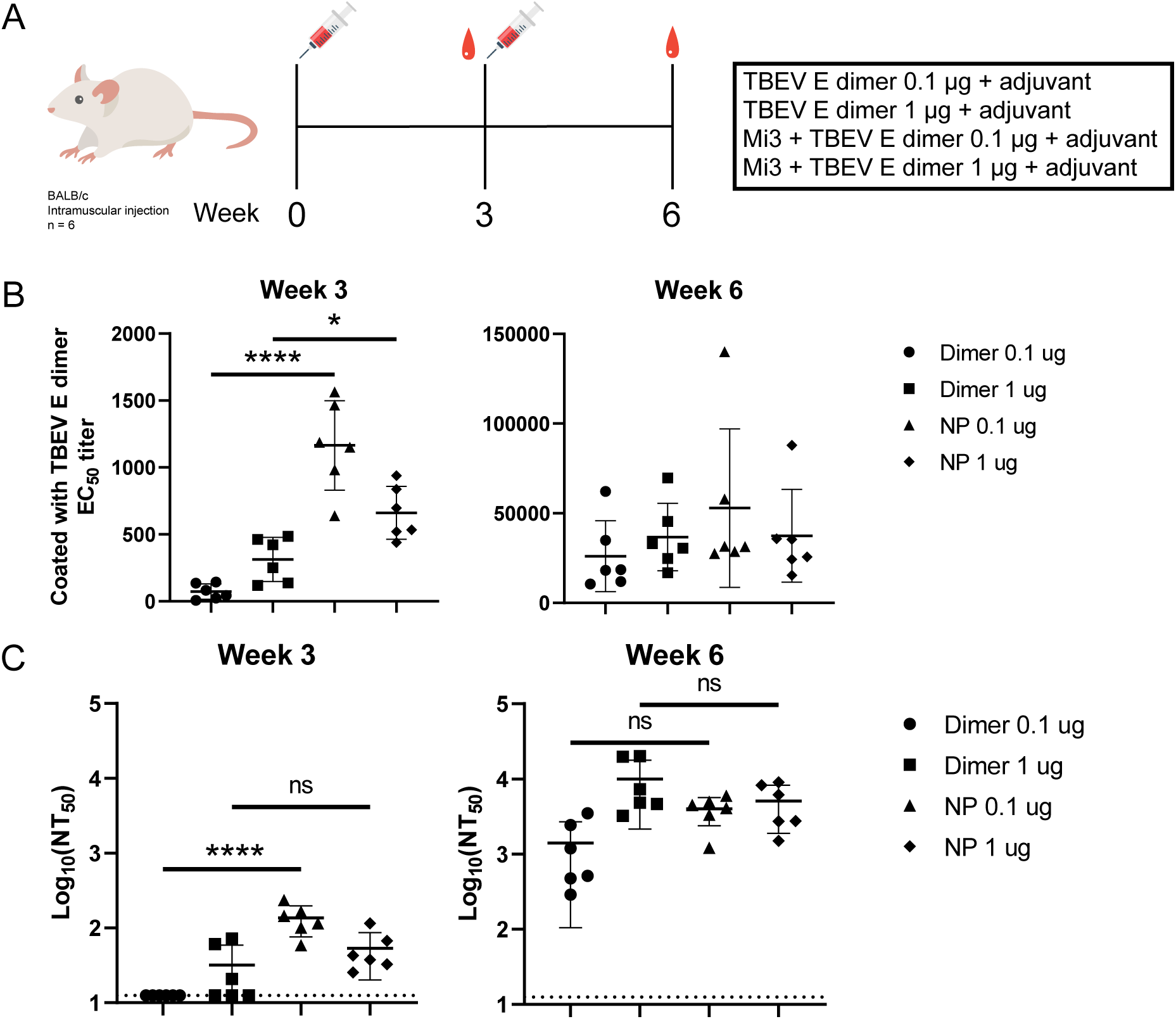
Immune Response of the TBEV E Dimer Nanoparticle Vaccine. (**A**) Immunization schedule for mice with the TBEV E dimer nanoparticle vaccine.(**B**) The IgG titers of mouse sera collected at each time point against TBEV E dimer were detected by ELISA. (**C**) The neutralization activities of sera after immunization with TBEV E dimer and nanoparticle vaccines against TBEV WH2012 were determined by the reporter virus particle neutralization assay. ELISA and the reporter virus particle neutralization assay were independently performed twice in duplicate wells. The representative data presented, as well as the half-maximal effective concentration (EC50) for ELISA and the half-maximal inhibitory concentration (NT50) for neutralization, were determined using nonlinear regression analysis. The dotted line indicates the limit of detection (LOD). Values below the LOD were calculated as the LOD. The data in each group were presented as the geometric mean and geometric standard deviation in one independent experiment. Statistical significance was determined by one-way ANOVA. ns indicates no significant difference, *p < 0.05, **p < 0.01, ***p < 0.001, ****p < 0.0001.

## DISCUSSION

In recent years, climate change and other factors have expanded the distribution range of ticks, leading to an increase in the incidence and diversity of tick-borne diseases[60]. In Europe, the number of cases of tick-borne encephalitis (TBE) is rising annually[61, 62]. Although inactivated virus vaccines currently approved can effectively reduce TBE cases, they require multiple doses for complete protection. Incomplete or irregular vaccination may still result in breakthrough infections[63]. Thus, there is a significant need for new vaccine formulations against TBEV to enhance control and eradication efforts. This study presents a TBEV E dimer nanoparticle vaccine that induces a robust humoral immune response in mice, highlighting its potential for further development and application.

The E protein of flaviviruses exists as dimers on the viral surface[64]. Evidence suggests that antibodies targeting epitopes formed between monomers within these dimers offer strong protection[65]. In this study, we introduced a cysteine mutation in TBEV E to create disulfide bond between interacting monomers, thereby forming stable dimers. Serum from mice immunized with the TBEV E dimer demonstrated significantly improved neutralization compared to serum from mice immunized with monomeric protein. This indicates that our TBEV E dimer effectively mimics the natural arrangement of E proteins on the virion surface, partially shielding non-neutralizing epitopes and enhancing the focus of humoral immunity on effective neutralizing sites.

Nanoparticle vaccines, which more closely resemble the size of viruses and present antigens more densely, can elicit stronger immune responses compared to recombinant protein subunit vaccines[66]. Nanoparticle vaccines have been utilized against various pathogens, including SARS-CoV-2[67], influenza virus[68], NIV[69], and SFTSV[70], with several demonstrating robust protective effects in preclinical models. In this study, Mi3 nanoparticles were employed to display the TBEV E dimer on their surface. The TBEV E dimer nanoparticle vaccine leveraged the advantages of the TBEV E dimer protein to induce enhanced humoral immunity, offering promising protection.

The expansion of tick distribution has led to the emergence of new TBEV subtypes through recombination. In addition to the traditional three subtypes, two additional TBEV subtypes have been identified. The TBEV E dimer not only exhibited excellent neutralization of the RVP of the homologous TBEV subtype but also effectively neutralized two additional subtypes.

In summary, we successfully designed and assembled the TBEV E dimer into a nanoparticle vaccine form. This TBEV E dimer nanoparticle vaccine induces a strong humoral immune response, focusing on neutralizing epitopes and effectively neutralizing three TBEV subtypes, demonstrating its broad-spectrum efficacy. The TBEV E dimer nanoparticle vaccine shows potential as a candidate for TBE prevention and control, and the dimer-based approach could be applicable to other flavivirus vaccines.

## MATERIALS AND METHODS

### Cell Culture

Vero E6, 293T, and K562 cells were maintained in DMEM (Monad) supplemented with 10% heat-inactivated FBS (ExCell Bio) at 37 °C and 5% *CO*_2_. Expi 293 cells were cultured in 293F Hi-exp medium (Opmbiosciences) under the same conditions.

### Plasmids and Cloning

Cloning was performed using Phanta Max Super-Fidelity DNA Polymerase (Vazyme) and the ClonExpress II One Step Cloning Kit (Vazyme). All open reading frames were validated by Sanger sequencing.

The sequences required for constructing the expression vectors included: pET28a-SpyCatcher003-Mi3 (GenBank MT945417), TBEV Far East subtype WH2012 (TBEV WH2012, GenBank KJ755186), TBEV European subtype Neudoerfl (TBEV Neu, GenBank TEU27495), TBEV Siberian subtype Vasilchenko (TBEV Vas, GenBank AF069066), ZIKV structural protein sequence (GenBank AQS26817), and DENV type I structural protein sequence (GenBank AAF59976.1).

The constructs His-SpyCatcher-Mi3, TBEV ED123, TBEVED12, TBEV ED3, TBEV ED123 4mFL and ZIKV ED12 were cloned into the pET21a prokaryotic expression vector. After successful sequencing verification, these constructs were transformed into BL21 (DE3) cells and induced with IPTG. His-SpyCatcher-Mi3 was expressed in soluble form, while other vectors were all obtained by inclusion bodies refolding.

The constructs TBEV prME-His, TBEV prME-D259C-His, TBEV prME-G262C-His, TBEV prME-V263C-His, TBEV prME-V263C-SpyTag-His, TBEV CprME-WH2012, TBEV CprME-Neudoerfl, TBEV CprME-Vasilchenko, ZIKV CprME, and DENV CprME were cloned into pCAGGS expression vectors. After successful sequencing and verification, the plasmids were purified for subsequent experiments.

### Expression and Purification of SpyCatcher-Mi3

The plasmid pET21a-His-SpyCatcher-Mi3 was transformed into E. coli BL21 (DE3) cells and plated on an ampicillin-resistant LB agar plate overnight at 37 °C. A single colony was then inoculated into 10 mL of LB medium containing ampicillin and incubated at 37 °C with shaking at 220 rpm for 16 hours. This culture was subsequently transferred to 1 L of LB medium containing ampicillin and grown at 37 °C with shaking at 220 rpm until the optical density (OD) reached 0.6–0.8. Protein expression was induced with 0.5 mM isopropyl *β*-D-1-thiogalactopyranoside (IPTG). The bacteria were then grown at 22 °C with shaking at 220 rpm for 16 hours. The culture was harvested by centrifugation at 8000 × g.

The cell pellet was resuspended in 100 mL of 250 mM Tris-HCl, 300 mM NaCl, pH 8.5, supplemented with lysozyme, DNase, *MgCl*_2_, and phenylmethylsulfonyl fluoride (PMSF). The lysate was incubated at room temperature for 2 hours. Ultrasonic disruption was performed on ice using an ultrasonic processor. The lysate was centrifuged at 8000 × g at 4 °C for 15 minutes to remove cell debris. The supernatant was filtered through a 0.45 *µ*m filter and purified using Ni-charge resin (GenScript). Endotoxins were removed by rinsing with a washing buffer containing 0.1% Triton X-114, followed by a rinse with a washing buffer without Triton X-114[71]. The protein was eluted with a washing buffer containing 300 mM imidazole. The purified protein was concentrated using a 100 kDa molecular weight cutoff centrifugal concentrator and stored at -80 °C.

### Mammalian Protein Expression

For mammalian protein expression, the plasmid was transfected into Expi293 cells using PEI (Polysciences). After 16 hours at 37 °C, the culture was transferred to 27 °C. The supernatant was harvested 108 hours post-transfection. The supernatant was filtered through a 0.45 *µ*m filter and the recombinant protein was purified using Ni-charge resin (GenScript). Further purification was achieved by size exclusion chromatography (SEC) using either a Superdex 200 Increase 10/300 GL column (Cyti-va) or a HiLoad 16/600 Superdex 200 pg column (Cytiva).

### Immunogen Preparation

The antigen was mixed with Mi3 in TBS at various molar ratios and incubated at 4 °C for 48 hours. Separately, uncoupled TBEV E dimer and uncoupled Mi3 were incubated at the same concentration in TBS at 4 °C for 48 hours. The optimal molar ratio for assembly was determined using SDS-PAGE/Coomassie staining. Subsequent assembly for immunization was carried out according to this optimized ratio.

### Negative Stain Electron Microscopy

The assembled nanoparticles were diluted to final concentrations of 800, 400, 200, 100, and 50 nM. A 10 *µ*L aliquot of each sample was adsorbed onto a 300-mesh copper grid (Beijing Daji Keyi Technology Co., Ltd., D11023) for 1 minute, followed by removal of excess solution with filter paper. The grid was stained with 2% uranyl acetate for 30 seconds, and the excess solution was blotted away. Images were acquired using a transmission electron microscope (JEOL, JEM-1400plus).

### Animal Experiments

BALB/c mice (6-8 weeks old, female) were obtained from Vital River Laboratory in Beijing. For the TBEV E dimer immunogenicity evaluation, the mice were randomly divided into three groups of six animals each. Mice were inoculated with 10 *µ*g of TBEV E dimer or 10 *µ*g of TBEV E monomer, while control mice were immunized with PBS. All groups were administered AddaVax adjuvant (InvivoGen) via intramuscular injection (i.m.) at 3-week intervals. Serum samples were collected 2 weeks after the final immunization, heat-inactivated at 56 °C for 30 minutes, and stored at -80 °C.

For the immunogenicity evaluation of NP-TBEV E dimer, BALB/c mice were divided into 4 groups, with six animals in each group. Mice were inoculated with either 0.1 or 1 *µ*g of TBEV E dimer - NP or TBEV E dimer. All groups received AddaVax adjuvant (InvivoGen) via intramuscular injection (i.m.) at 3-week intervals. Serum samples were collected 3 weeks after each immunization, heat-inactivated at 56 °C for 30 minutes, and stored at -80 °C.

### Mouse Antiserum ELISA

Purified protein was immobilized on a 96-well high-binding flat-bottom microplate (Corning) at 200 ng per well in carbonate buffer and incubated at 4 °C overnight. After washing the wells three times with PBST (PBS containing 0.05% Tween 20), the plate was blocked with PBS containing 5% skim milk powder (MPBS). Diluted mouse antiserum was added and incubated at room temperature for 1 hour. Following three washes, HRP-conjugated anti-mouse IgG (H + L) diluted in 5% MPBS (1:8000, ABclonal) was added and incubated for 1 hour. After additional washing, TMB (3,3’,5,5’-tetramethylbenzidine) substrate was applied, and the reaction was stopped with 1 M HCl. Absorbance was measured at 450 nm. Data were analyzed using GraphPad Prism 8.

### Reporter Virus Particle (RVP) Production

Following the manufacturer’s instructions, GeneTwin (Biomed; TG101) was used for co-transfection of 1 *µ*g pWNVI-GFP-replicon plasmid and 3 *µ*g of the selected flavivirus CprME plasmid into 293T cells to produce RVP. After a 5-hour incubation at 37°C, the medium was replaced with DMEM containing 10% heat-inactivated FBS. Supernatants containing RVP were collected every 24 hours, filtered through a 0.45 *µ*m filter, and frozen at -80°C. The medium was refreshed with DMEM containing 10% heat-inactivated FBS. The frozen RVP was subsequently titrated on Vero E6 or 293T cells.

### RVP Neutralization Test

RVP, which can produce 400 to 600 fluorescent spots, was mixed with 100 *µ*L of diluted serum and incubated at 37°C for 1 hour. Subsequently, 80 *µ*L of the mixture was added in duplicate to a 96-well plate with Vero E6 or 293T cells (2000 cells per well). After incubation at 37°C for 48 hours, cells were fixed with 4% paraformaldehyde, and green fluorescent dots (GFP signals) were counted using a CTL-S6 Universal M2. The semi-maximum suppression concentration (NT50) values were determined using a nonlinear regression model and plotted with GraphPad Prism 8.

### Antibody-Dependent Enhancement Assay

A 100 *µ*l aliquot of diluted RVP was mixed with 100 *µ*l of diluted serum and incubated at 37°C for 1 hour. Subsequently, 80 *µ*l of the mixture was added in duplicate to a 96-well plate containing K562 cells (2000 cells per well). The plate was incubated at 37°C for 48 hours, after which the cells were fixed with 4% paraformaldehyde. The percentage of cells exhibiting green fluorescence was analyzed using flow cytometry with the CytoFLEX S (Beckman Coulter). The Fold change was calculated by dividing the positive percentage of the experimental group by the positive percentage of the blank group without serum.

### Antigen-Specific IgG Depletion Assay

To remove antigen-specific IgG antibodies, the protein with Strep tag was incubated with Streptactin Beads (Smart-Lifesciences; SA092005) for binding. Each round included equilibrating 20 *µ*l of microspheres in the binding buffer (PBS or DMEM) and coupling them with 5 *µ*g of recombinant protein at 4°C for at least 45 minutes. Unbound proteins were removed by centrifugation, and the beads were washed with PBS before adding the diluted serum for specific antibody depletion. This process was performed for six rounds to ensure complete depletion. Subsequently, ELISA and RVP neutralization experiments were conducted.

### Statistical Analysis

All data were analyzed and graphs were generated using GraphPad Prism 8, and the values were expressed as the mean ± standard deviation (SD). One-way or two-way analysis of variance (ANOVA) was performed between the experimental groups, and a p-value less than 0.05 indicated a statistically significant difference. Ns indicates no significant difference (ns, p > 0.05; *, p < 0.05; **, p < 0.01; ***, p < 0.001; ****, p < 0.0001).

## CONFLICTS OF INTEREST

The authors declare no conflict of interest.

## Supplementary Figures

**FIG S1.**
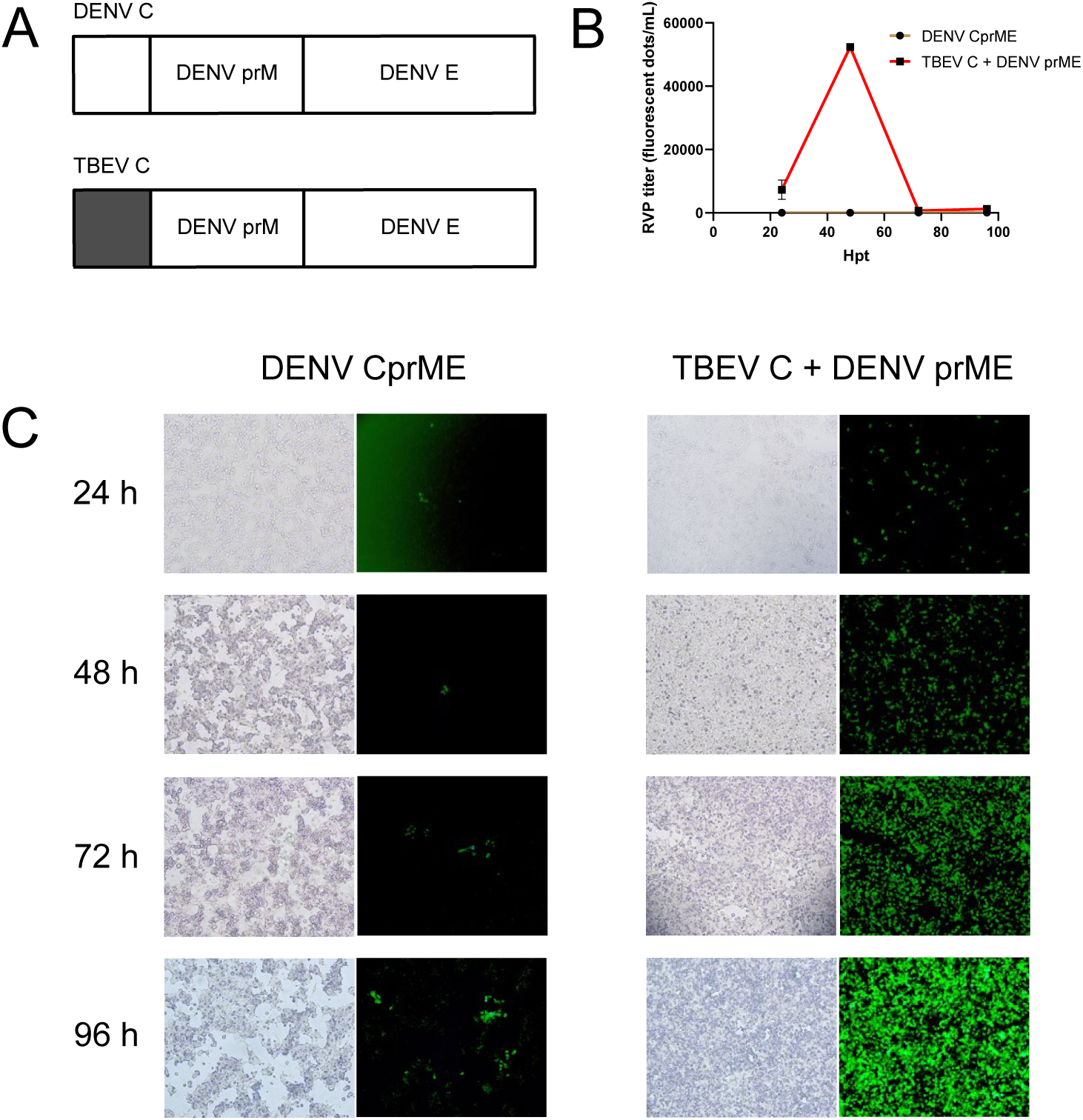
Preparation of RVP DENV. (**A**) A schematic diagram of the DENV structural protein CprME. We replaced the C of DENV with TBEV C. (**B**) After co-transfecting TBEV C - DENV prME and replicon, supernatants were collected every 24 hours, and the supernatants were subjected to titer determination of RVP DENV. (**C**) After co-transfecting TBEV C - DENV prME and replicon, the growth of green fluorescence in cells was observed every 24 hours.

**FIG S2.**
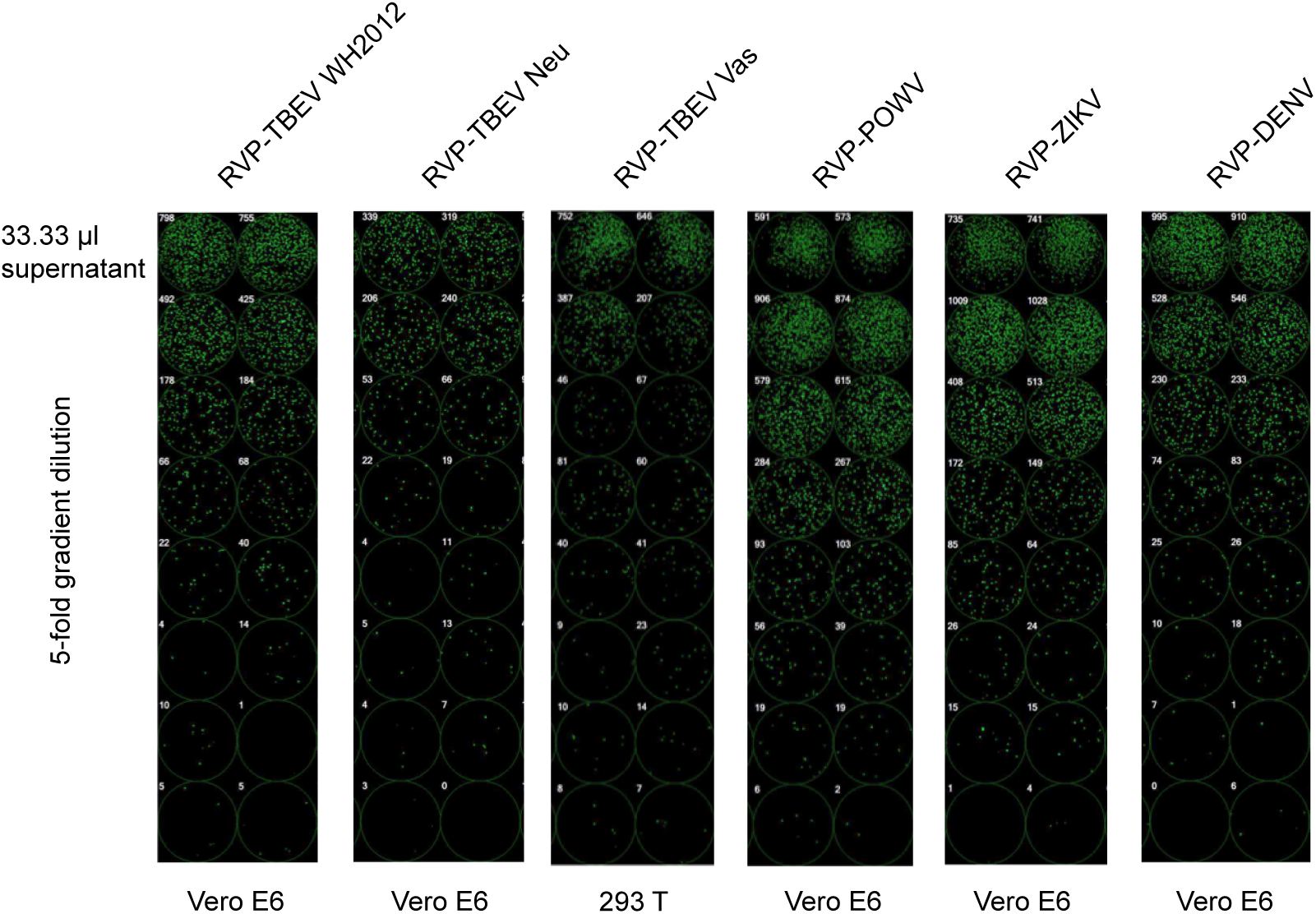
Titration of RVPs of different flaviviruses. This study involved a total of 4 flaviviruses, namely TBEV, POWV, ZIKV and DENV. Among them, three subtypes of RVPs of TBEV were prepared: RVP - TBEV WH2012, RVP - TBEV Neu and RVP - TBEV Vas.

**FIG S3.**
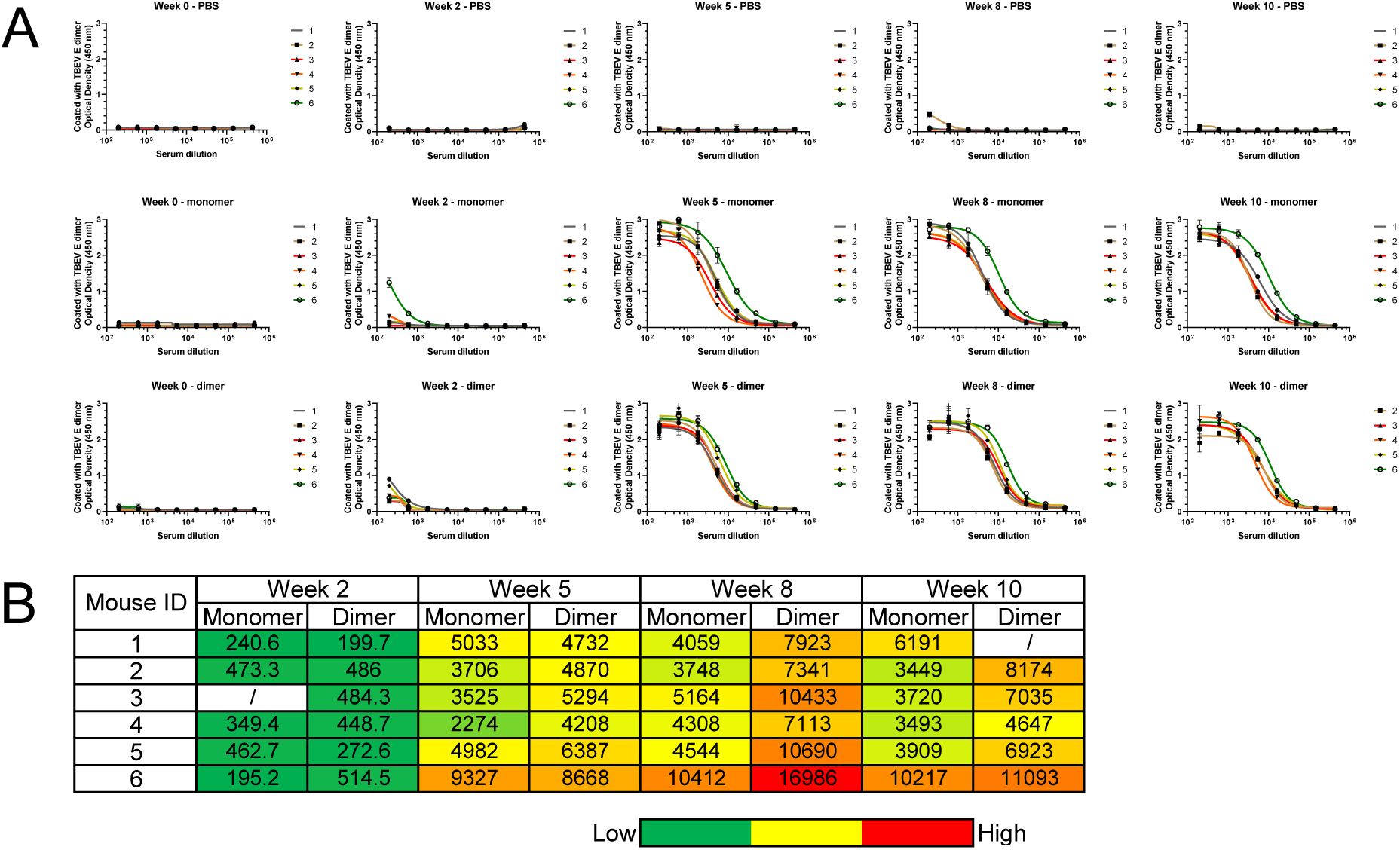
The IgG titers of mouse sera binding to the TBEV E dimer after immunization with the TBEV E dimer and monomer proteins. (**A**) The ELISA binding curves of specific IgG antibodies against TBEV E dimer in the serum of each mouse in the three immunization groups at different time points. The time points for blood collection were Week 0, Week 2, Week 5, Week 8, and Week 10. (**B**) Summary of EC50 titers measured for three immunization groups against the TBEV E dimer coating antigen. Color coding represents the level of EC50 titer (from green to red, indicating low to high). It’s important to note that the serum binding of the control group and the sera at Week 2 of the experimental groups did not reach a plateau (or saturation), which makes it challenging to accurately determine the EC50 titers. Nevertheless, the EC50 values calculated in GraphPad Prism 8 were used as a quantitative measure of binding antibody titers to facilitate the comparison of different groups. The experiments were independently conducted twice in duplicate, and all the data are represented as mean ± standard error.

**FIG S4.**
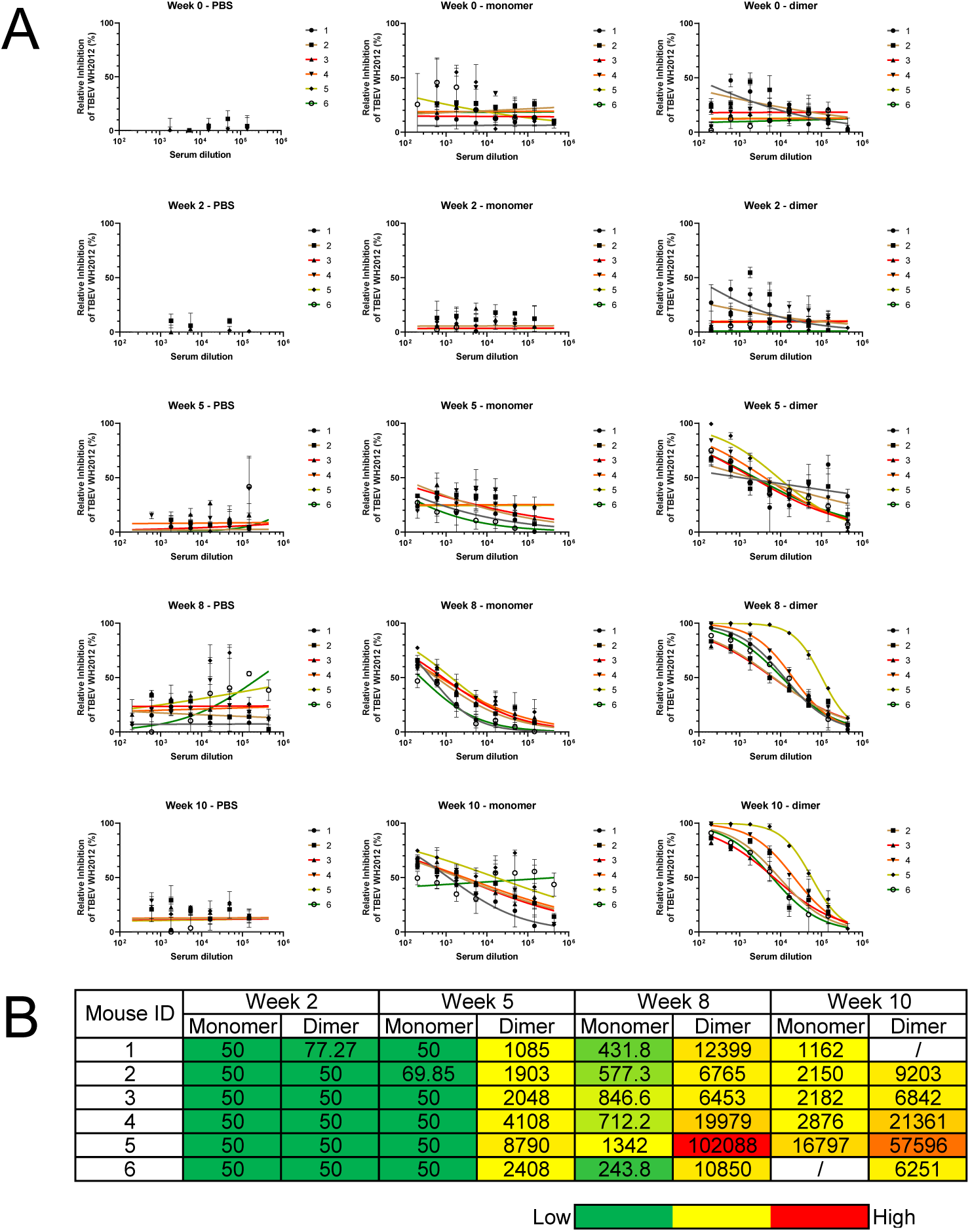
The titers of neutralizing antibodies in mouse sera from different immunization groups at different time points. (**A**) The inhibition curves of each mouse serum in the three immunization groups against RVP - TBEV WH2012 at different time points. The time points for blood collection were Week 0, Week 2, Week 5, Week 8, and Week 10. (**B**) Summary of IC50 titers measured for three immunization groups against RVP - TBEV WH2012. Color coding represents the level of IC50 titer (from green to red, indicating low to high). It’s important to note that the serum binding of the control group and the sera of the experimental groups in the first 2 to 3 weeks did not reach a plateau (or saturation), which makes it challenging to accurately determine the IC50 titers. Nevertheless, the IC50 values calculated in GraphPad Prism 8 were used as a quantitative measure of neutralizing antibody titers to facilitate the comparison of different groups. The experiments were independently conducted twice in duplicate, and all the data are represented as mean ± standard error.

**FIG S5.**
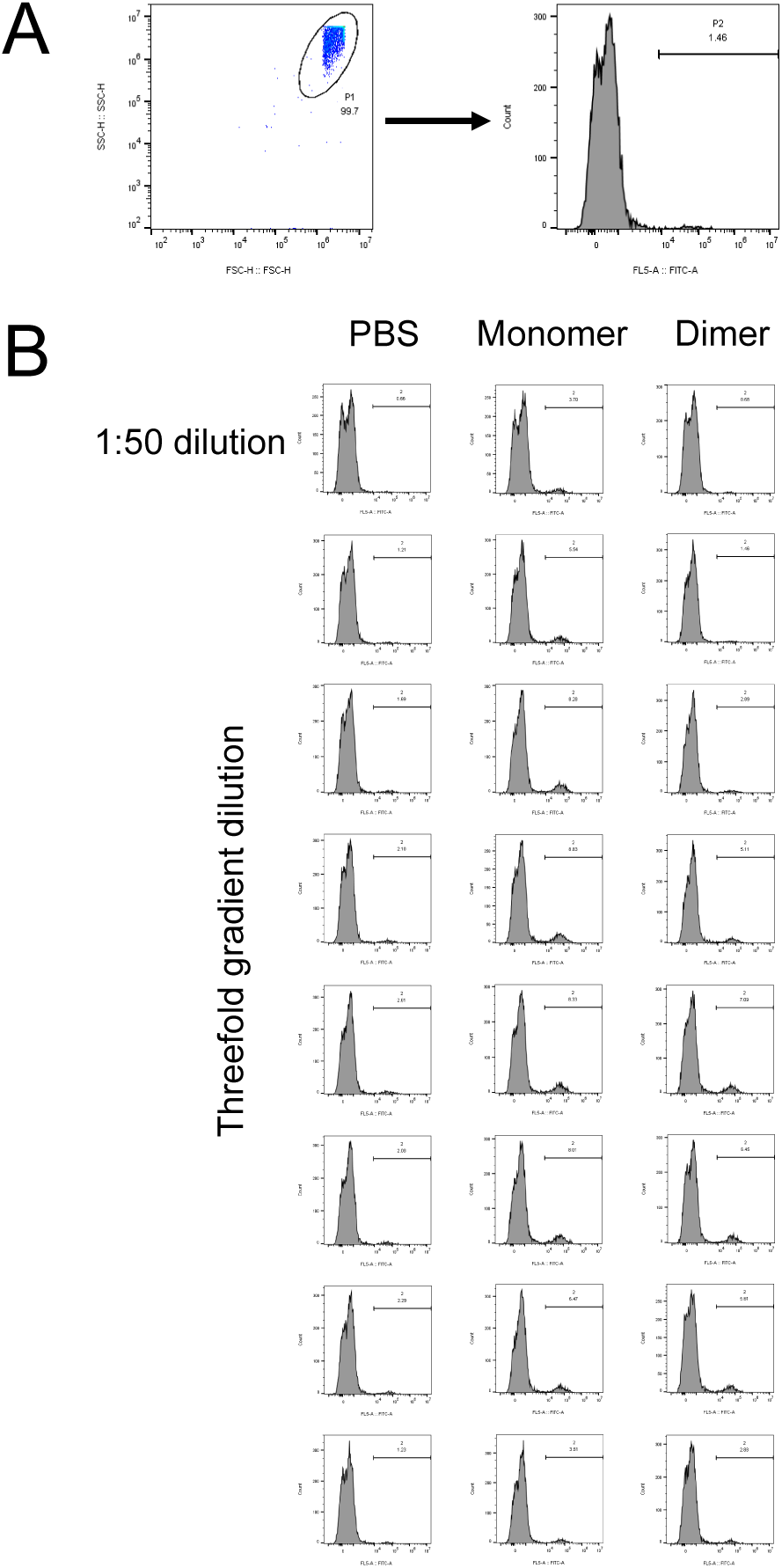
Analysis of antibody-dependent enhancement of infection experiments (ADE). (**A**) The gating strategy of flow cytometry. After RVP infection, it will express green fluorescence, corresponding to FITC. (**B**) The ADE effect of the sera from three mice in the three immunization groups after immunization on RVP WH2012.

**FIG S6.**
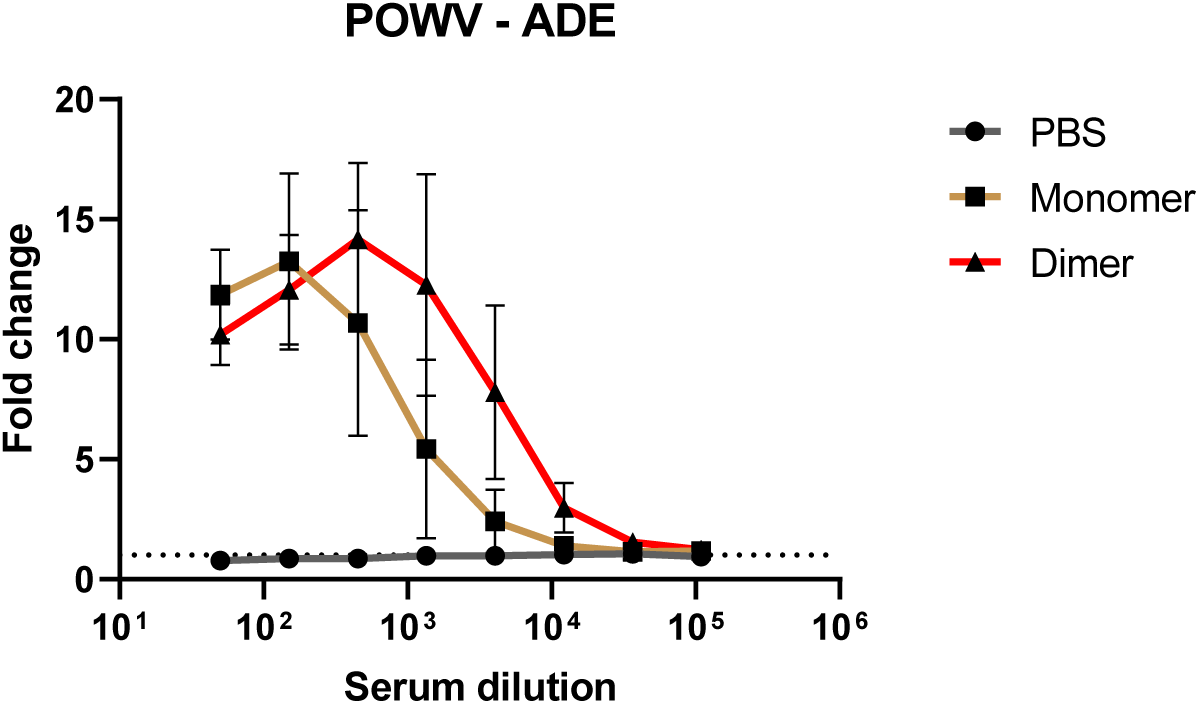
The ADE effect of mouse sera at Week 8 in the three immunization groups on RVP POWV.

**FIG S7.**
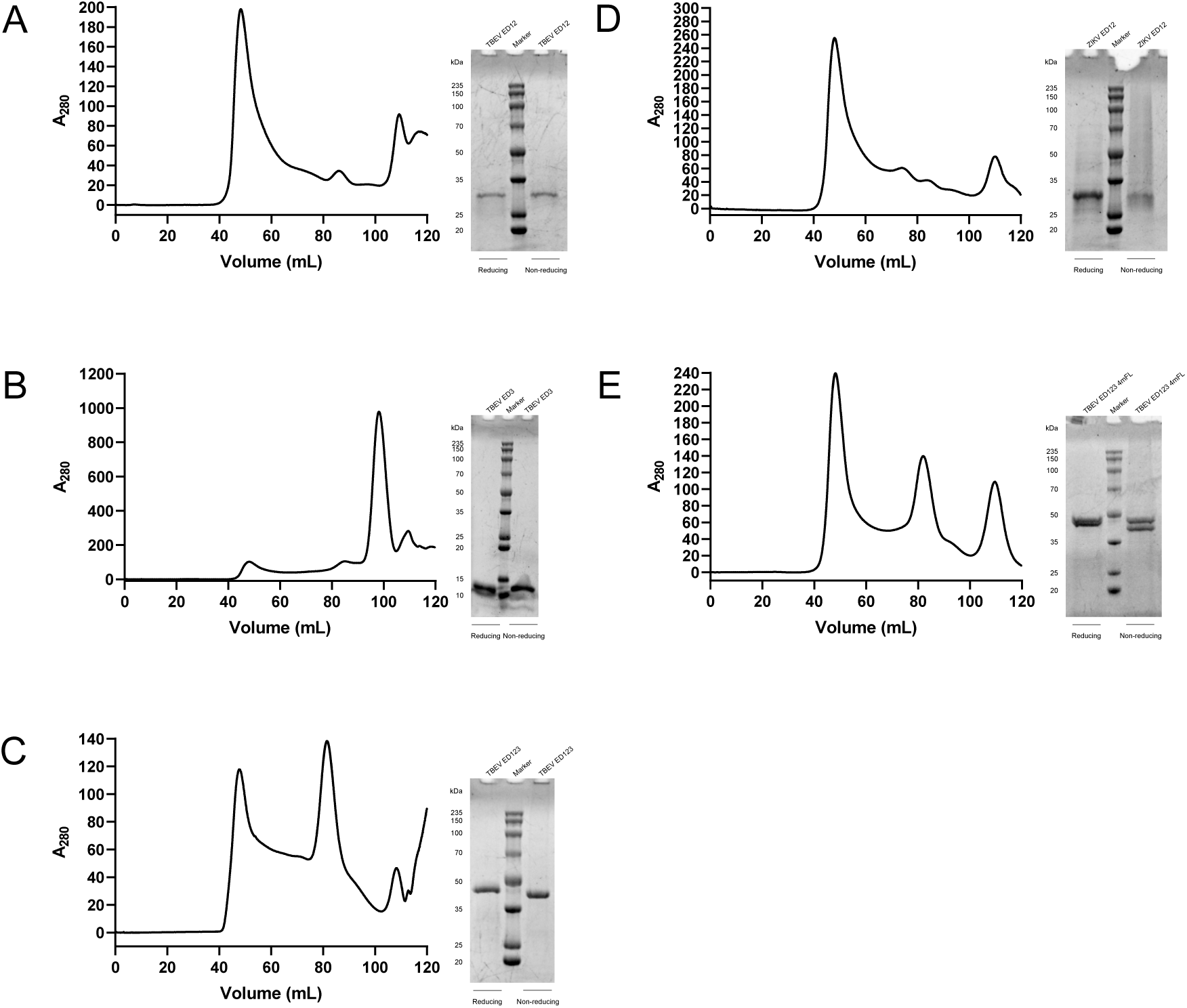
Expression and purification of different domain proteins of TBEV and ZIKV. Here are shown their size-exclusion chromatography (SEC) and SDS-PAGE/Coomassie staining results respectively. (**A**) TBEV ED12.The peak position is approximately at 85 mL. (**B**) TBEV ED3.The peak position is approximately at 101 mL. (**C**) TBEV ED123.The peak position is approximately at 83 mL. (**D**) ZIKV ED123.The peak position is approximately at 84 mL. (**E**) TBEV ED123 4mFL.The peak position is approximately at 83 mL.

**FIG S8.**
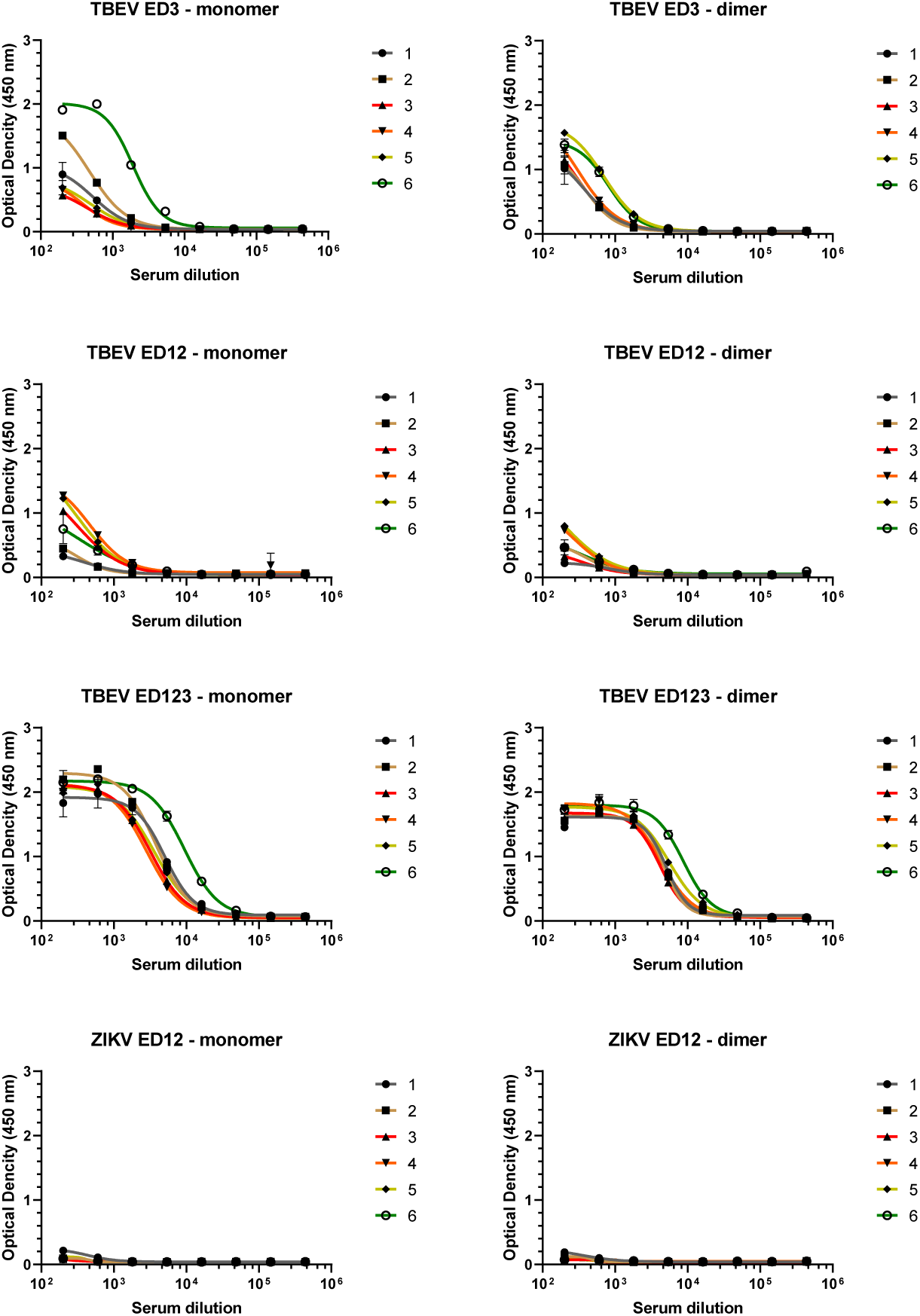
The binding curves of sera from each mouse in the dimer and monomer protein immunization groups against TBEV ED3, ED12, ED123 and ZIKV ED12 at Week 5.

**FIG S9.**
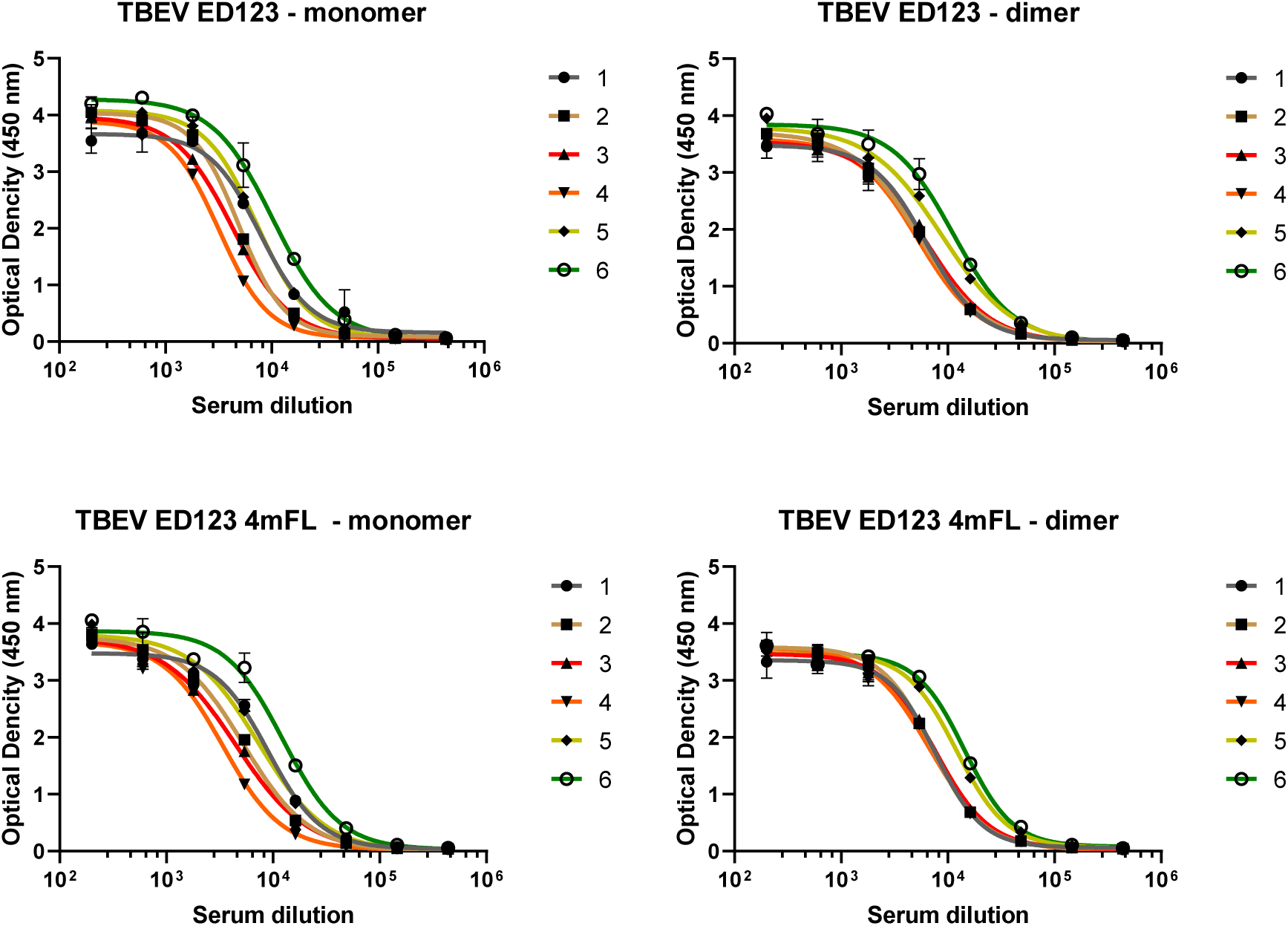
The binding curves of sera from each mouse in the dimer and monomer protein immunization groups against TBEV ED123 and TBEV ED123 4mFL at Week 5.

**FIG S10.**
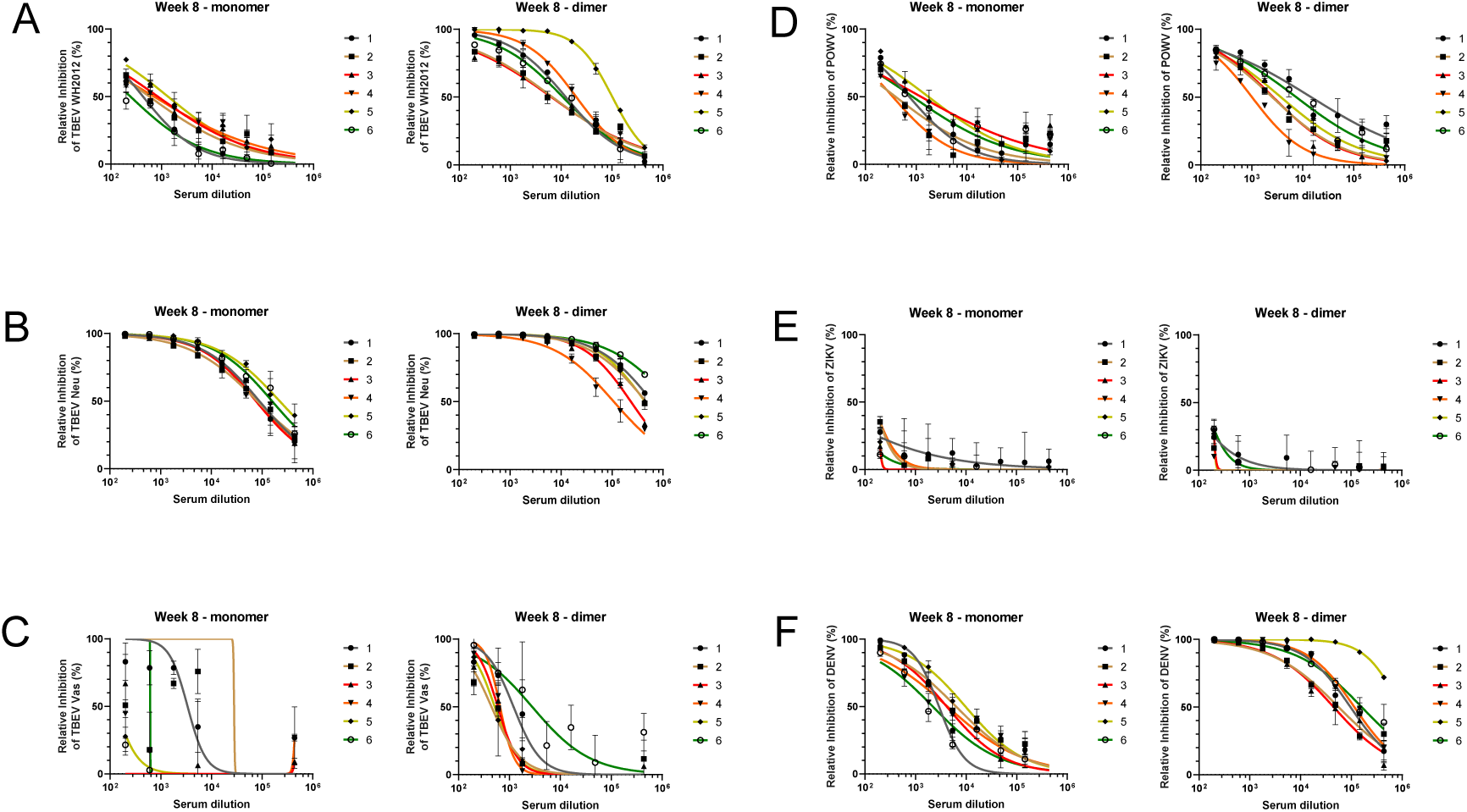
The inhibition curves of sera from each mouse in the dimer and monomer protein immunization groups against different flaviviruses at Week 8. (**A**) RVP - TBEV WH2012. (**B**) RVP - TBEV Neu. (**C**) RVP - TBEV Vas. (**D**) RVP - POWV. (**E**) RVP - ZIKV. (**F**) RVP - DENV.

**FIG S11.**
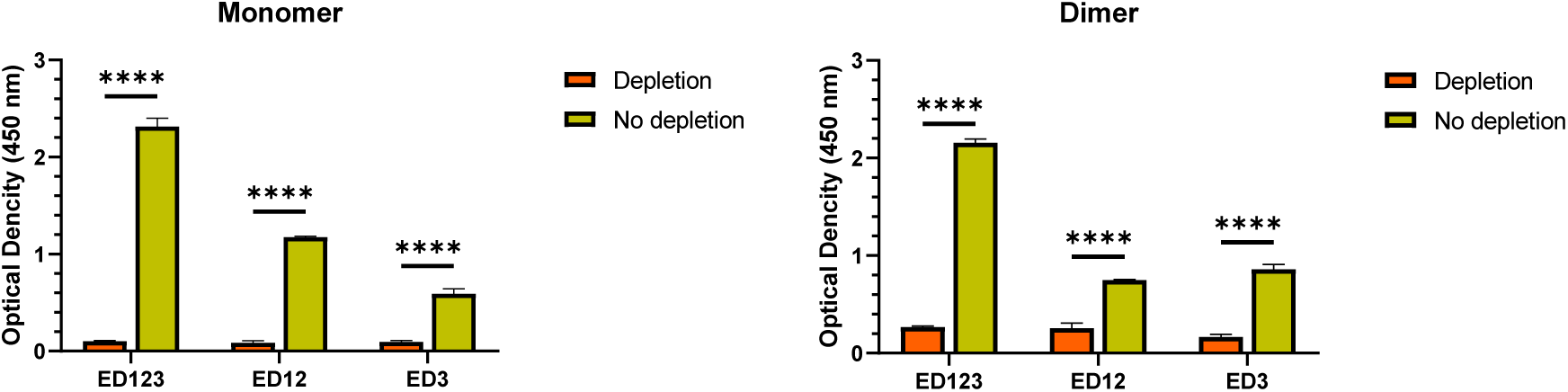
After the antibody depletion experiment, ELISA was used to verify the depletion effect. On the left is the binding intensity of the immune serum of the monomer protein to the protein of the domain after depletion of the protein of different domains, and on the right is that of the immune serum of the dimer protein.

**FIG S12.**
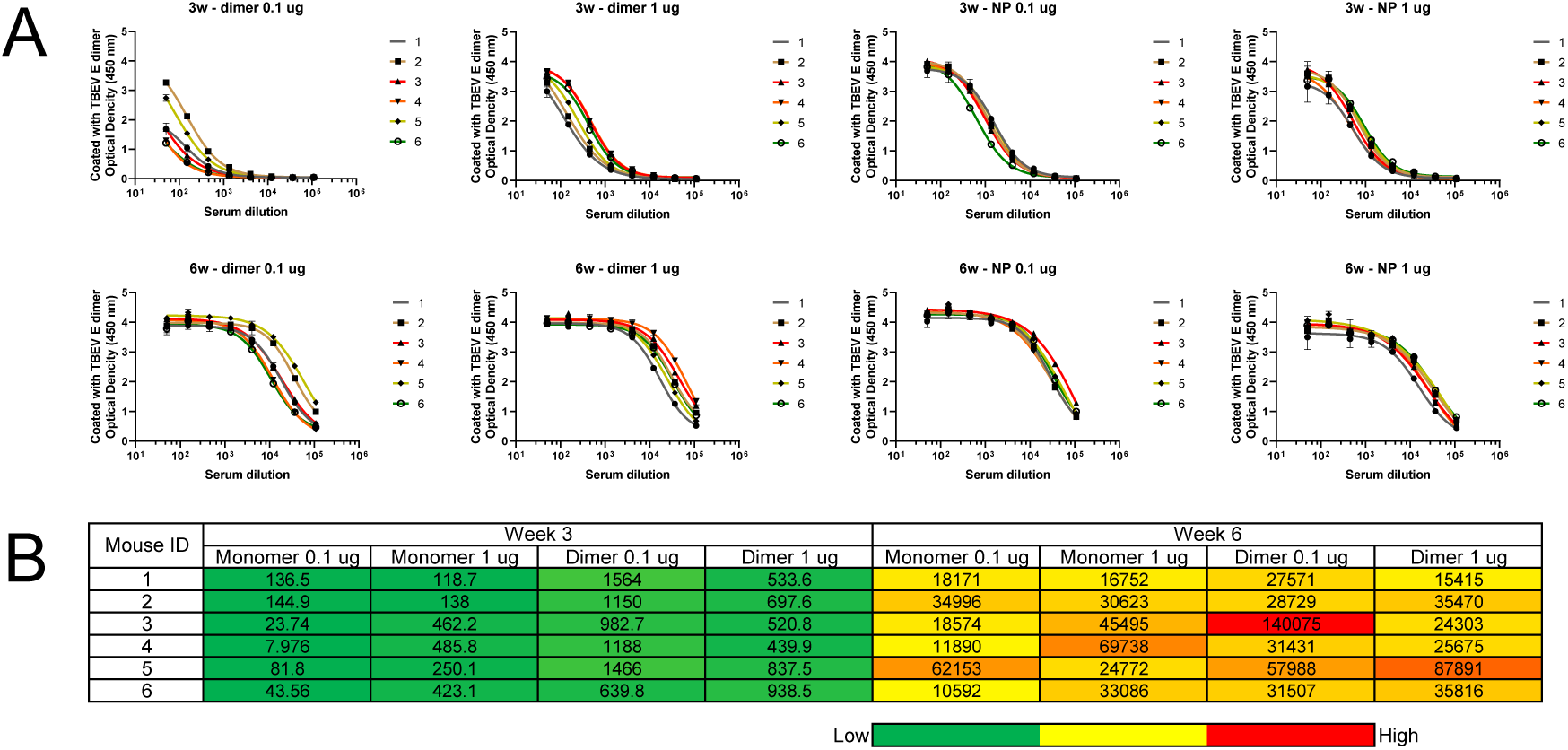
The IgG titers of mouse sera binding to the TBEV E dimer after immunization with the TBEV E dimer and dimer nanoparticles. (**A**) The ELISA binding curves of specific IgG antibodies against TBEV E dimer in the serum of each mouse in the four immunization groups at different time points. The time points for blood collection were Week 3 and Week 6. (**B**) Summary of EC50 titers measured for the four immunization groups against the TBEV E dimer coating antigen. Color coding indicates the level of EC50 titer (from green to red, representing from low to high). It should be noted that the serum binding of the experimental group in Week 3 did not reach a plateau (or saturation), which makes it challenging to accurately determine the EC50 titers. Nevertheless, the EC50 values calculated in GraphPad Prism 8 were used as a quantitative measure of binding antibody titers to facilitate the comparison of different groups. The experiments were independently conducted twice in duplicate, and all data are presented as mean ± standard error.

**FIG S13.**
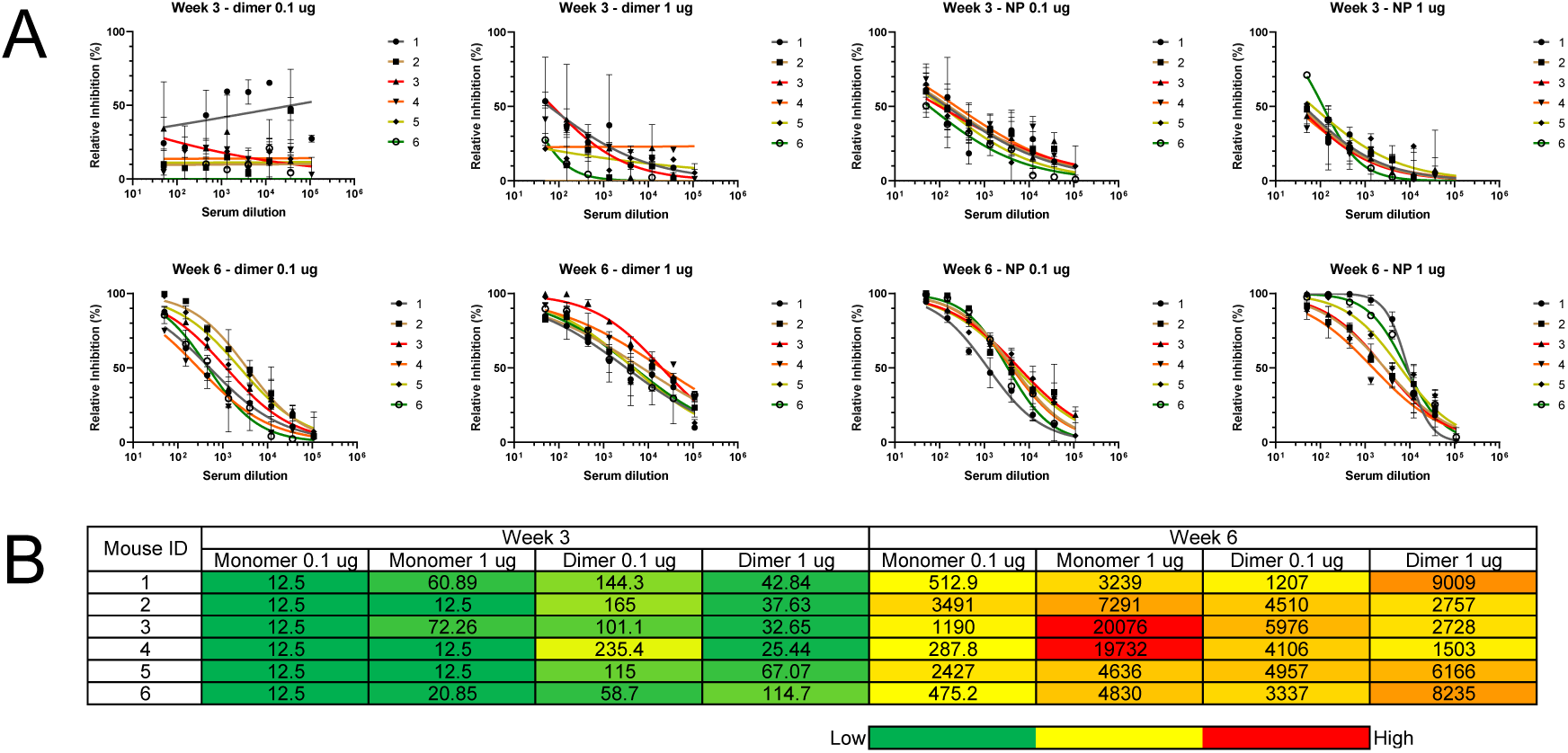
The titers of neutralizing antibodies in mouse sera from different immunization groups at different time points. (**A**) The inhibition curves of sera from each mouse in the four immunization groups against RVP - TBEV WH2012 at different time points. The time points for blood collection were Week 3 and Week 6. (**B**) Summary of IC50 titers measured for the four immunization groups against RVP - TBEV WH2012. Color coding represents the level of EC50 titer (from green to red, indicating low to high). It’s important to note that the serum binding of the experimental group at Week 3 did not reach a plateau (or saturation), which makes it challenging to accurately determine the IC50 titers. Nevertheless, the IC50 values calculated in GraphPad Prism 8 were used as a quantitative measure of neutralizing antibody titers to facilitate the comparison of different groups. The experiments were independently conducted twice in duplicate, and all the data are represented as mean ± standard error.

## Notes

### Competing Interest Statement

The authors have declared no competing interest.

